# Harmonising digitised herbarium data to enhance biodiversity knowledge: creating an updated checklist for the flora of Greenland

**DOI:** 10.1101/2024.12.01.626242

**Authors:** Brandon Samuel Whitley, Jakob Abermann, Inger Greve Alsos, Elisabeth M. Biersma, Viktor Gårdman, Toke Thomas Høye, Laura Jones, Nora Meriem Khelidj, Zhao Li, Gianalberto Losapio, Thomas Pape, Katrine Raundrup, Paula Schmitz, Tiago Silva, Helena Wirta, Tomas Roslin, Natalie Iwanycki Ahlstrand, Natasha de Vere

**Author notes:** Corresponding Author: Natasha de Vere, Natural History Museum of Denmark, University of Copenhagen.

## Abstract

International efforts to digitise herbarium specimens provide the building blocks for a global digital herbarium. However, taxonomic changes and errors can result in inconsistencies when amalgamating specimen metadata, that compromise the assignment of occurrence records to correct taxa, and the subsequent interpretation of patterns in biodiversity. We present a novel workflow to mass-curate digital specimens. By employing existing digital taxonomic backbones, we aggregate specimen names by their accepted name and flag remaining cases for manual review. We then validate names using site-specific floras, balancing automation with taxonomic expert-based curation. Applying our workflow to the vascular plants of Greenland, we harmonised 175,266 digitised herbarium specimens and observations from 92 data providers from the Global Biodiversity Information Facility (GBIF). The harmonised metacollection for the Greenland flora contains 780 plant species. Our workflow increases the number of species known from Greenland compared to other currently available species checklists and increases the mean number of occurrences per species by 42.6. Our workflow illustrates the integration required in order to create a global, universally accessible digital herbarium, and shows how previous obstacles to database curation can be overcome through a combination of automation and expert curation. From the specific perspective of the Greenland flora, our approach arrives at a new checklist of taxa, a new curated metacollection of occurrence data, and revised estimates of plant richness. The list of taxa and their prevalence allow a new basis for biodiversity assessment and conservation planning.

**Societal Impact Statement:** Digitising plant collections has allowed for data to be aggregated across multiple collections, forming a single harmonised resource of unprecedented scale. This resource is only accurate once the database names are assigned to one accepted name per species. We established a semi-automated workflow for processing plant name data, leveraging taxonomic backbones and employing taxonomic expertise at key stages. Applying our workflow to the flora of Greenland, we developed a curated checklist of 780 species, capturing greater species richness than previously published, while also curating 175,266 plant records. Our findings redefine our knowledge of Greenlandic plant diversity, while harmonising a vast digital collection for further research.

## Introduction

The rapid adoption of widespread herbarium digitisation, coupled with other forms of digital data acquisition (such as citizen science observations), has produced an unprecedented volume of publicly available data on the occurrence of species over space and time. By uploading these data to public data infrastructures such as the Global Biodiversity Information Facility (GBIF) (http://www.gbif.org), previously separate collections become accessible and amalgamable into a single metacollection, ultimately laying the foundation for a global digital herbarium (Davis, 2023).

While improving over time, there is still a considerable lack of occurrence data on many plant species for many habitats (Feeley, 2015; Vargas et al., 2024). Fieldwork is often positioned as a key solution to filling gaps in data (Vargas et al., 2024). However, the nomenclatural and geographical curation of existing biological occurrence data has been suggested to be a more effective first-step (Vargas et al., 2024). Such curation has been shown to deflate estimates of alpha diversity, while increasing accuracy. At the same time, the spatial coverage of species increases, as synonymous entries are aggregated (Vargas et al., 2024).

At the forefront of this curation process is the need to harmonise the taxonomy and nomenclature of these collections, aggregating specimens locally curated under various synonyms and with variant name spellings or author accreditations by a single accepted taxon name (Groom et al., 2019; Maldonado et al., 2015). There are many ways in which incongruencies can occur when processing taxonomic and nomenclatural data in digital collections. Such incongruencies can result in a mismatch of names, resulting in an improperly reconciled metacollection. Incongruencies primarily occur within five categories: variants in spelling, transcription errors, taxonomic changes, missing authorship, and errors in name recognition.

Variants in taxon name spelling or author spelling and format can hinder data harmonisation, such as *Ligusticum scothicum* L. being spelled as *Ligusticum scoticum* L.; or *Potentilla rubella* T.J.Sørensen being authored as *Potentilla rubella* Sørensen. Such variants can result in inflated estimations of diversity. Cardoso et al., (2017) found that 7% of the lowland Amazon rain forest tree species listed in an Amazonian tree flora were synonyms or spelling variants. Thus, taxonomic expertise is required to validate whether the names denote the same taxa, as minor differences in spelling cannot be consistently handled through an automated process. To illustrate, most automated fuzzy match processes would incorrectly join *Carex disperma* Dewey to *Carex trisperma* Dewey, which are two different species.

While there are resources available to process some of these incongruencies, such as the Taxonomic Name Resolution Service (TNRS) (Boyle et al., 2021), automated nomenclatural decision processes can result in incorrect harmonisation depending on the parameters selected. For example, GBIF specimen NHMD814312, comes with the interpreted name (i.e. the name which GBIF has ascribed to it from one of its many taxonomic backbones) of *Aira alpina* Roth, 1789. Upon inspecting the digitised herbarium sheet, this specimen can have its name verified as *Aira alpina* L.. Yet, the best match offered by TNRS is *Aira alpina* Sobol. ex Rupr., a synonym of *Avenella flexuosa* subsp. *flexuosa*. Thus, while resources such as the TNRS are undoubtedly powerful, the need for manual curation checks is still emphasised by studies using their services (Wenk et al., 2024). The similarity of spelling for some different taxon names, and the availability of multiple correction options for spelling errors, contribute to the imperfect success of major nomenclature checking software. Consequently, manual expertise cannot be replaced with complete automation when dealing with inconsistencies and nuances (Wagner, 2016).

Errors in the transcription of data from digitised specimens can also lead to harmonisation errors, whereby the label may specify the name of one species, whereas what is uploaded into GBIF is the name of another. This is especially common in cases of older herbarium labels, which can be difficult for humans and artificial intelligence alike to transcribe.

An accepted name is the name in current use for a given taxon, for which all other names under which it is known become synonyms. As taxonomic revisions occur within a given group the name under which an occurrence is labelled can change from accepted name to synonym and can therefore be missed when querying for an accepted name without accounting for its synonyms. Incongruencies can also occur when differentiation information is missing or incorrect, such as the author information for a name. A species name alone is not always enough to distinguish it, as there are cases of different species having an identical name but different authors. For instance, *Arenaria caespitosa* J.Vahl is a synonym of *Sagina caespitosa* Lange, while *Arenaria caespitosa* Phil. is a synonym of *Arenaria serpens* Kunth.

There can also be errors in recognising the name of a species, either due to the quality of the taxonomic backbone at hand, or due to the name not having been published or formally recognised by the broader scientific community. An example of this is the GBIF specimen NHMD1195336, labelled as *Luzula arctica* f. *pygmæa*, although this name is not recognised in major taxonomic backbones. These cases can often represent new species determinations worthy of further study, but in which there has been a lag between new determination, validation, and publication (Bebber et al., 2010).

Through the synthesis of a metacollection, the actual biodiversity contained within the collection emerges, optimising data available for further analysis, such as creating a species checklist for a given geographic area (Ball-Damerow et al., 2019; Victor et al., 2014). This has a broad range of applications, including selecting relevant species to include in DNA barcode reference libraries (Jones et al., 2021), biodiversity monitoring (Ball-Damerow et al., 2019), invasive species monitoring (Pagad et al., 2022; Reaser et al., 2020), conservation planning and status updating, designation of protected areas (Marín-Rodulfo et al., 2024) and leveraging large-scale biodiversity data to test theories in ecology and evolution (Ramirez-Parada et al., 2024).

Harmonising the metacollection by nomenclature and taxonomy allows for all possible occurrences linked to one accepted name to be brought together for analysis, maximising the data pool for a species in size, and by collection time, geographic location, collector, and collection context. This can be particularly important when working with species that have undergone major taxonomic revisions over time, such as historical specimens collected under the genus *Aira* which are now known to belong to *Avenella* and *Deschampsia*. By harmonizing taxa names, we can capture any cases of taxonomic change and enable access to data from more specimens. This optimises the available data for a broad range of species-specific ecological research applications (Lang et al., 2019; Meineke et al., 2018), including machine learning training (de Lutio et al., 2022; Pearson et al., 2020), distribution modelling, interaction, trait, and plasticity analyses (Heberling, 2022; Heberling & Isaac, 2017; Pearson et al., 2020; Willis et al., 2017), genomic based research (Alsos et al., 2020; Nevill et al., 2020), and conservation status research (Albani Rocchetti et al., 2021), while also informing future species collection endeavours (James et al., 2018). Expanded species-specific occurrences can also be used in humanities research, where a harmonised metacollection can expand the availability of data and associated metadata for use in research related to the history of science, ethnobotany, and colonial legacies (Groom et al., 2014; Ningthoujam et al., 2014; Park et al., 2023; van Andel et al., 2012).

Here, we introduce a novel workflow to mass harmonise digital nomenclatural and taxonomic data for collections to contribute to the development of a global digital herbarium. We aim to demonstrate an efficient, traceable, and iterative workflow for curating the nomenclatural and taxonomic data of a digital collection derived from multiple data providers and covering a multitude of names and naming variants for different species and taxa.

We use Greenland (Kalaallit Nunaat) as our case study. Along with the Arctic in general, parts of this region are warming at four times the global average (Rantanen et al., 2022). Greenland is experiencing large-scale changes in plant composition, plant phenology, and widespread shrubification (Grimes et al., 2024; Høye et al., 2013), making the study of its flora of critical importance to understanding and predicting further impacts of climate change. Greenland contains all Arctic bioclimatic zones, and thus a high percentage of its flora is also found in other Arctic and sub-Arctic biomes. This sharing of species among regions highlights the relevance of harmonising the nomenclatural data (Elven et al., 2017; Raynolds et al., 2019). The latest flora of Greenland was published in 1978 (Böcher et al., 1978), with an update on new records for species and distributions in 2020 giving 532 vascular plant species for Greenland (Bay, 2020). However, a recent large-scale digitisation project has contributed a high number of specimens (ca. 170,000 from 145,000 herbarium sheets) to the Greenland vascular plant digital collection available on GBIF (Iwanycki Ahlstrand, 2023) making a valuable resource that potentially contains more species digitally available for the first time.

Our workflow optimises the balance between automated processing and expert taxonomic review, producing a curated metacollection of GBIF occurrences with harmonised nomenclatural data. We then use these data, along with a reference flora, to produce a species (taxon) checklist for the flora of Greenland. Our resulting checklist therefore acts as a summary of historical occurrence data, utilising existing taxonomic decisions surrounding the metacollection while also providing an iterative process that can be continually updated through modern taxonomic decisions.

## Materials and Methods

A workflow was developed to download GBIF plant occurrence data and subject the nomenclature assignment of the occurrences to a semi-automated curation process (Figure 1). The resulting output is a curated Taxon List along with a curated metacollection of occurrences.

**Figure 1:**
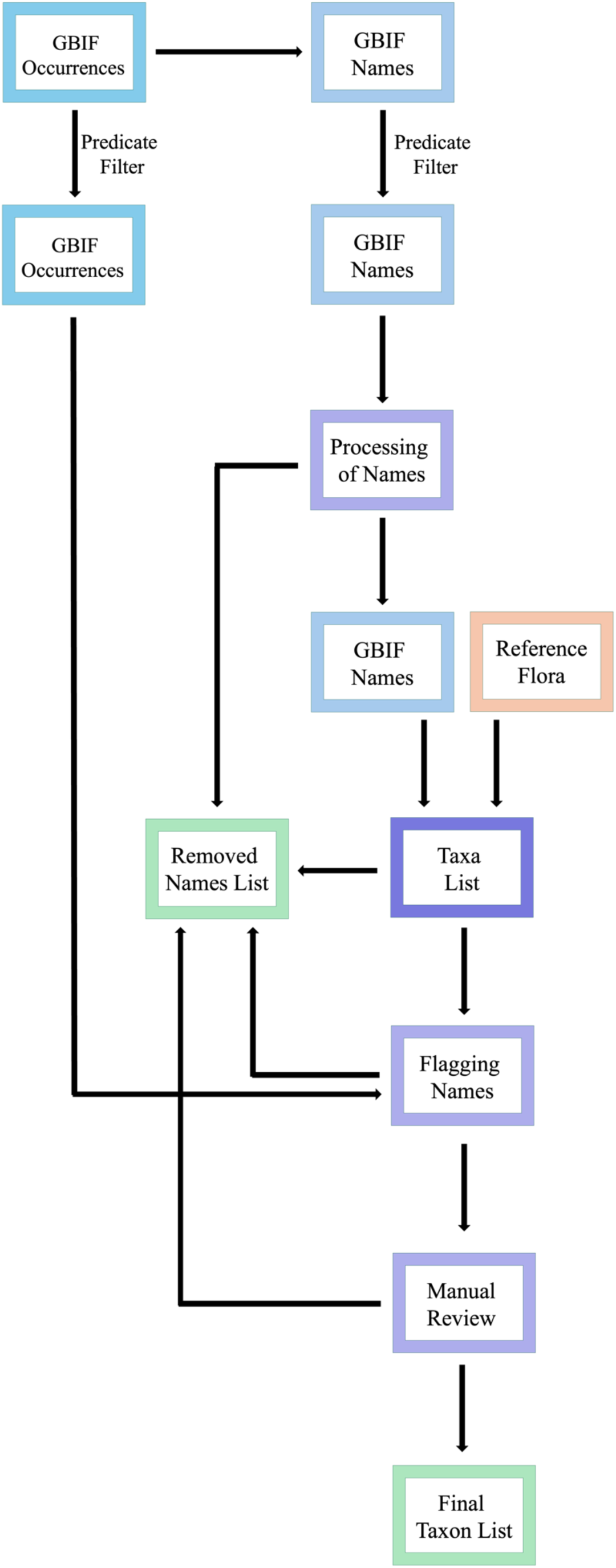
A simplified flowchart of the workflow used to curate a metacollection of GBIF vascular plant records. GBIF occurrences were downloaded, and the resulting unique occurrence names were extracted. These names underwent automated and manual processing and curation, resulting in some names being removed (stored in a Removed Names List), some names being corrected, and some not needing manual manipulation. Thereafter, an external reference flora for the geographic area was integrated into the Taxon List. In the next step, the names were matched back to the corresponding GBIF occurrences, and the number of occurrences and other metadata were used to flag remaining taxa names for review. Finally, the remaining taxa names were compared to external checklists, and species unique to the Taxon List were manually reviewed. A final curated Taxon List was produced, based on the GBIF occurrence data and that of the reference flora, along with a metacollection of Greenland vascular plant occurrence data. A full description of the workflow is described in Methods S1.

### Querying Data

GBIF occurrence data for the vascular plants of Greenland was downloaded (https://doi.org/10.15468/dl.yta9wk) into R (V4.2.2) using the *rgbif* package (Chamberlain, 2017; R Core Team, 2022). Filters specific to the GBIF platform were built into the workflow such that the user can specify their query (i.e. to specific plant groups, geographic regions, types of records). The records included both herbarium specimens and research-grade human observation records. Using the scientific names (assigned by GBIF to each occurrence), all unique names were collected for processing (Methods S1).

### Processing Names

To process the names, the *KEWR* package (Walker, 2023), was used to run names against the Plants of the World Online (POWO) database to determine their nomenclatural and taxonomic status (POWO, 2024). Query results indicated if the name was the accepted name for the taxon (to have an accepted nomenclatural status), if it was a known synonym to an accepted name, or if it was an unrecognised name. Unrecognised names required manual checking of associated occurrence data, largely composed of digitised herbarium specimens. Through this process, the unrecognised name could either be corrected to a known accepted name or synonym or maintained as unrecognised.

Names which did not come with an author were processed depending on if their name had matches to multiple authors (Methods S1). Autonym names (i.e. names which contain the epithet for the typical infraspecific taxon) were processed separately (Methods S1).

### Integration of an external reference flora

An external reference flora more localised to the geographic region of focus was selected. For our case study of Greenland, the Annotated Checklist of the Panarctic Flora Vascular plants (PAF) was used (Elven et al., 2017). This flora was integrated into the Taxon List (Methods S1). Remaining unrecognised names were first checked against the reference flora. Any unrecognised names still remaining were removed and placed into a removed names list (a list to track all names removed during the workflow at different stages).

### Internal validations – flagging names of concern

The GBIF occurrence data was appended with the final harmonised accepted name for each given occurrence. A single accepted name could now be queried across all the GBIF data, and all occurrences matching to that accepted name or any of its synonyms could be gathered.

Using the number of occurrences and occurrence metadata, names were flagged if they did not meet predefined criteria. Validation criteria were set such that any accepted name containing five occurrences or less (customizable), or any accepted name only based on human observation (and not digitised herbarium specimens) was flagged for review. Additionally, an option was coded to flag names based on a minimum number of contributing data sources (customisable but preset to NULL), to enable further customisation for other users. Flagged names were then checked through a semi-automated process to determine if they were to be kept in the Taxon List (Methods S1).

### Applying taxonomic expertise

Remaining flagged names were subjected to manual review using taxonomic expertise. For our case study of Greenland, we first unflagged names where the observation had been validated by known experts in the field and based on our knowledge of the local flora. We then unflagged names that were found in external databases of Greenlandic plants (Methods S1).

### Comparison to External Checklists

The remaining Taxon List was then compared to five external checklists for Greenland (Methods S1):

1. The Greenland Flora (1978, updated in 2020) – a checklist of native and naturalised taxa listed in the 1978 Grønlands Flora (Böcher et al., 1978), transcribed and updated in 2020 (Bay, 2020; Jacobsen et al., 2020).
2. The Wildflowers of Greenland – a field guide for the vascular plants of Greenland (Rune, 2011).
3. The KEW POWO taxa listed for Greenland – flora listed as occurring in Greenland in the KEW POWO database (POWO, 2024).
4. The Red List Assessment for Greenland’s Plant Species (Boertmann & Bay, 2018).
5. The Adventitious Plants and Cultivated Plants in Greenland – a checklist of introduced plants found in Greenland (Jacobsen, 2023; Pedersen, 1972).

Species from the Taxon List not found in the external checklists were selected for manual review and were validated using the images attached to the occurrences, knowledge of the flora, and individual research on each species (Methods S1).

### A Final Taxon List

After processing all flagged names, the final Taxon List was produced. Additional outputs of the process included a removed names list, containing all names that had been removed during the workflow, specimen counts data (number of occurrences and occurrence age range per accepted name or removed name). Additionally, two GBIF metacollection files were produced, one for the Taxon List and one for the removed names list, where every occurrence was assigned to an accepted name or a removed name.

### Consequences for estimates of biodiversity

To understand how the harmonisation of the data affects our emergent understanding of biodiversity within Greenland, we used the resulting database to derive Hill Numbers. Hill Numbers characterise the different facets of community-level diversity, where q0 describes taxa richness (i.e. independent of taxon occurrence abundance), q1 describes the exponential of Shannon’s entropy index (i.e. diversity when taxa are weighted in proportion to their frequencies), and q2 describes the inverse of Simpson’s concentration index (i.e. diversity when abundant taxa are proportionally weighted more than rarer taxa) (Chao et al., 2014; Jost, 2006; Roswell et al., 2021). By comparing diversity across these orders, we can examine the changes in effective sample size, thus capturing a broader range of biodiversity properties. Given the unique qualities of our dataset (containing reference flora taxa names with no occurrences), we chose to also calculate taxa richness directly and compare changes across taxonomic ranks and harmonisation steps using a Chi-squared test.

## Results

### An updated checklist of Greenlandic vascular plants

Our filtered GBIF query resulted in 175,266 occurrences from 92 data sources and corresponded to 3674 unique taxon names. The harmonisation workflow resulted in fewer taxon names compared to the starting list from the GBIF data (Figure 2). Our resulting Taxon List for Greenland contains a total of 1239 accepted names, corresponding to 167,263 (95.4%) of the GBIF occurrences (Figure S1). This included 18 families, 150 genera, 1 section, 733 species, 232 subspecies, 67 varieties, 1 form, 21 hybrids, and 16 species aggregates (Table 1). When adding names that only occur at the infraspecific level to the species level, our Taxon List contains a total of 780 vascular plant species for Greenland, a 29% and 61% increase in number of species as compared to the 603 reported by the combined Greenland Flora of Böcher et al. (1978), Bay (2020) and Jacobsen et al. (2020), and the 483 species reported in the Wildflowers of Greenland (Rune, 2011). Our Taxon List contained 76 species not found on any of the external checklists (including 56 found exclusively from the GBIF occurrence data), making up nearly 10% of the species in our Taxon List. The full workflow outputs can be found in Dataset S1 and Dataset S2.

**Figure 2:**
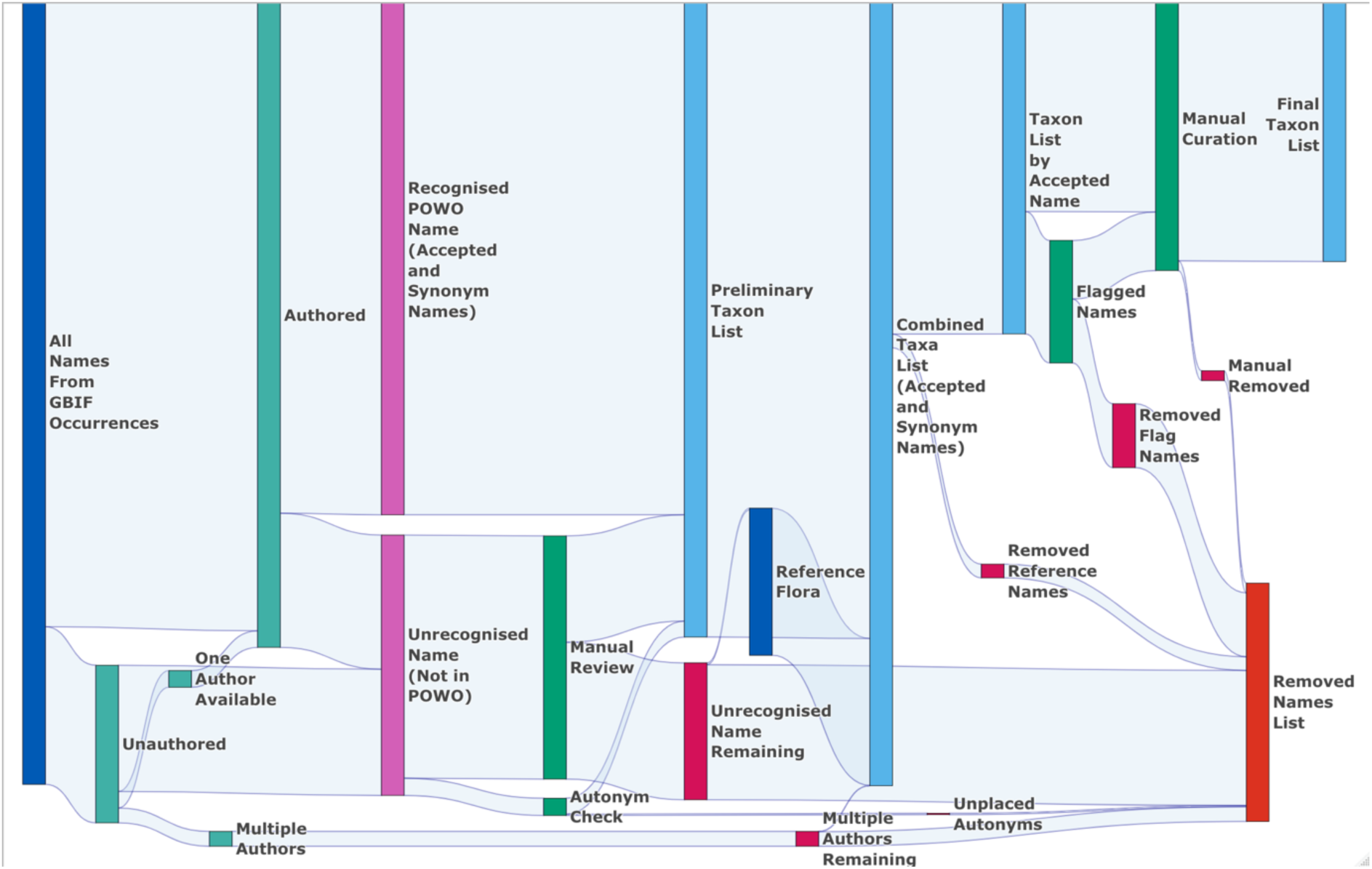
Sankey diagram summarising the full workflow process of creating the metacollection of Greenlandic vascular plants from GBIF occurrence data and an integrated reference library. 3674 unique taxon names provided by GBIF were allocated to 1239 Accepted Names, corresponding to 167,263 (95.4%) of the GBIF occurrences. These occurrences can now be treated as a single metacollection. A full description of the workflow is described in Methods S1.

**Table 1:** Summary of the biodiversity found within our Taxon List, partitioned by taxon rank, compared to five external lists of Greenlandic plant species. For each external checklist, and for all external checklists combined, the total number of taxa per species, subspecies, and variety, and the percent which match to our Taxon List is shown. * When bringing infraspecific names to a species level, our Taxon List contains a total of 780 vascular plant species. Data sources include 1. The Greenland Flora (containing native and introduced species) (Bay, 2020; Böcher et al., 1978; Jacobsen et al., 2020), 2. The Wildflowers of Greenland (a field guide for the vascular plants of Greenland) (Rune, 2011), 3. The KEW POWO flora listed for Greenland – flora listed as occurring in Greenland in the KEW POWO database (POWO, 2024), 4. The Red List Assessment for Greenland’s Plant Species (containing native species) (Boertmann & Bay, 2018), 5. The Adventitious plants and cultivated plants of Greenland (a checklist of introduced plants) (Jacobsen, 2023; Pedersen, 1972).

|  | **Species** |  | **Subspecies** |  | **Variety** |  |
| --- | --- | --- | --- | --- | --- | --- |
| Our Taxon List | 733* |  | 232 |  | 67 |  |
|  | **Total** | **% Match** | **Total** | **% Match** | **Total** | **% Match** |
| Greenland Flora<sup>1</sup> | 557 | 97.5% | 77 | 96% | 25 | 48% |
| Wildflowers of Greenland<sup>2</sup> | 457 | 99.3% | 51 | 98% | 6 | 100% |
| KEW Plants of the World: Greenland<sup>3</sup> | 599 | 98.5% | 168 | 100% | 34 | 100% |
| Red List<sup>4</sup> | 431 | 98% | 40 | 92.5% | 9 | 100% |
| Adventitious Plants<sup>5</sup> | 156 | 83.3% | 6 | 66% | 1 | 0% |
| Combined External Lists | 746 | 94.5% | 207 | 96.6% | 50 | 74% |

### Comparison to external checklists reveals high correspondence

Upon cross referencing our final Taxon List with the five external lists, we found a high correspondence of matching taxa. Of the 1050 unique names of species rank or lower found when combining all five external lists, only 73 (7%) were not found in our Taxon List (Dataset S1). Of these unmatched names, 41 are species, 7 are subspecies, 13 are varieties, 10 are forms, and 2 are hybrids, and 16 of these unmatched names are shared among one or more external lists (Table 1). At a species level, this corresponded to 45 species from the external lists being not found in our Taxon List. The external list containing adventitious and cultivated plants in Greenland (Jacobsen, 2023; Pedersen, 1972) had the least correspondence to our Taxon List across species, subspecies, and varieties.

### Harmonising nomenclature decreases alpha diversity but increases occurrences per accepted name

Alpha diversity (taxon richness regardless of number of occurrences) of ranks under genus level significantly decreased for all taxon ranks (except hybrids) through harmonisation (Table S1). In all ranks except hybrids, there were significant differences in overall alpha diversity between the original GBIF data and the two stages of the harmonisation process (Table S2). The integration of the Panarctic reference flora resulted in no significant change in alpha diversity across any ranks below genus level, indicating high correspondence of reference flora names to the existing names sourced from GBIF occurrences.

By harmonising the nomenclatural and taxonomic data, the effective sample size of each taxon decreased for total taxon richness (q0), abundance and evenness (q1), and dominance and evenness (q2) (Figure 3). Thus, the harmonisation of the Greenland vascular plant nomenclature data not only reduced total taxon richness, but the remaining data became less evenly distributed (in terms of occurrences per taxon). Fewer taxa are therefore contributing more to occurrence composition. Notably, even before nomenclatural harmonisation, there is clear evidence of unevenness and dominance in the occurrence data, suggesting that some taxa compose larger proportions of the collection, with many taxa being rare. While this property was expected, it was increased by harmonisation, where rarer taxa names were aggregated to common taxa.

**Figure 3:**
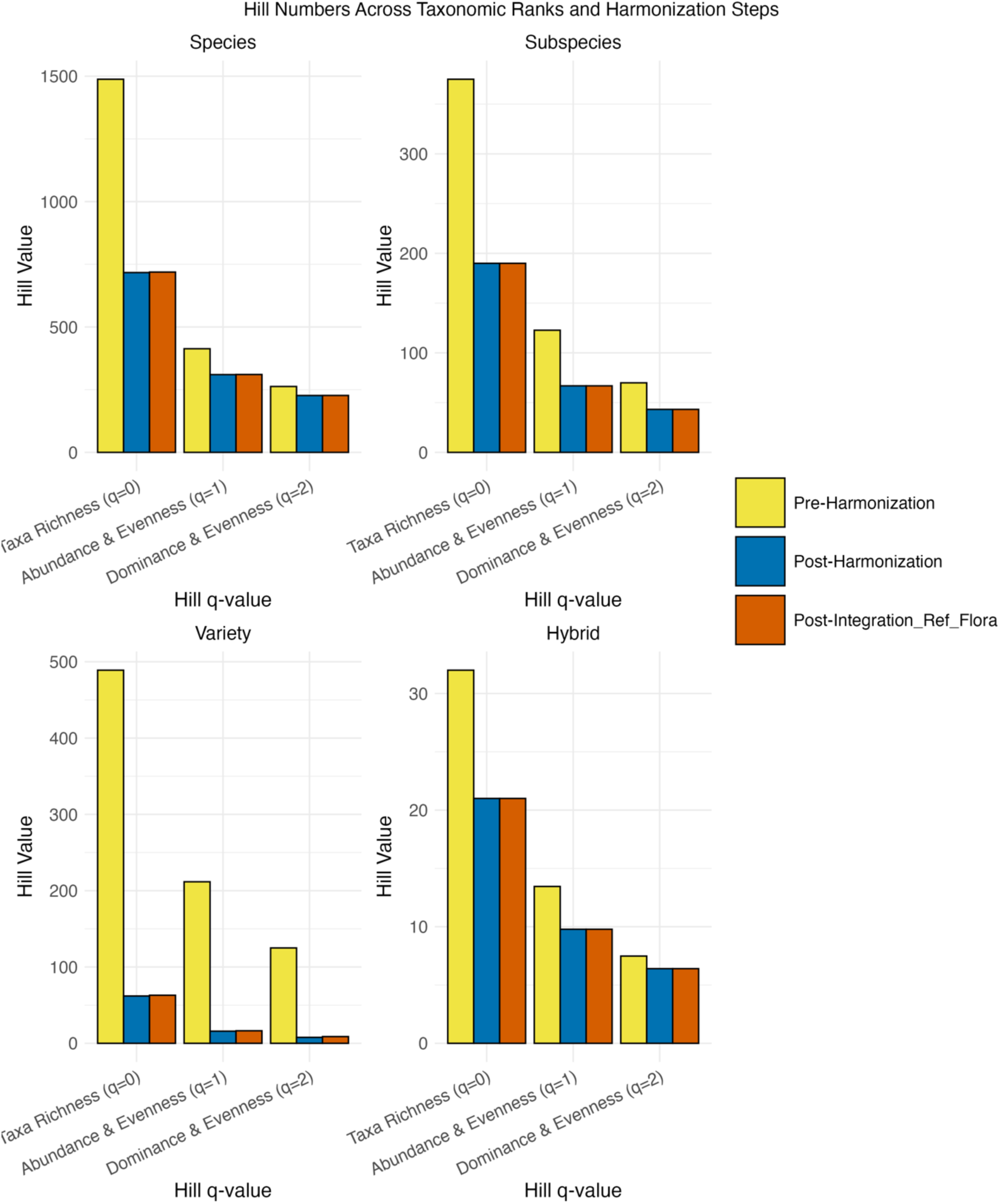
Bar plots divided by taxon rank, indicating the Hill Values (effective number of taxa or species equivalents) on the Y-axis, and the diversity order (ranging from 0-2) on the X-axis. For each diversity order, the Hill Values are shown for the GBIF occurrence data before harmonisation, after harmonisation, and after integrating the Panarctic reference flora. Forms were excluded due to low data count.

The harmonisation process significantly increased the number of occurrences per accepted name, with a mean change of 42.6 new occurrences and the mean time series of the accepted names by 15.7 years (Figure S2). This further supports the differences seen in Figure 3, whereby specific taxa are becoming more dominant as a result of data harmonisation.

### Removed Names

1052 names were removed from the Taxon List due to at least one of nine possible reasons (Table 2). Of these, unrecognised names formed the majority of the removed names.

**Table 2:** Summary of the removed names list, indicating the number of names which have been removed per removal category.

| Reason for Name Removal | Number of Names |
| --- | --- |
| Unrecognised name | 556 |
| Specimen metadata (incorrect sampling location, corrupt image) | 127 |
| Accepted name with low number of occurrences (customised criteria) | 125 |
| Unauthored name with multiple author options | 66 |
| Invalid name integrity (name spelling could not be properly discerned) | 64 |
| Reference list name with multiple POWO matches | 62 |
| Manually removed (removed in final curation stage) | 44 |
| Autonym name with multiple author options for parent species | 6 |
| Unplaced name with matching accepted name or synonym | 2 |
| Total | 1052 |

Our harmonisation process resulted in 556 unrecognised names, divided into 161 varieties, 140 forms, 90 species, 85 subspecies, 76 hybrids, 2 genera or higher, and 2 sections. The genera with the most unrecognised names included *Ranunculus* (59 names), *Draba* (51 names), and *Carex* (39 names). In some cases, herbarium sheets have been revised from their original designation to a new one, and the new designation is not known to the POWO database or the reference flora. As an example, 45 out of 59 names in the genus *Ranunculus* have been redetermined to unrecognised names by two researchers in the 1980s. Furthermore, in the majority of cases the unrecognised name is an infraspecific designation to a known species name, but the infraspecific designation itself was not known to the taxonomic databases.

## Discussion

### A species checklist for the vascular plants of Greenland

Through our workflow, we were able to generate a representative checklist for the vascular plants of Greenland, making use of the considerable amount of GBIF occurrence data available, while corroborating with a reference flora. Through this harmonisation process, the GBIF data used has been curated for its nomenclature, resulting in a derived GBIF dataset which can now function as a single metacollection. This offers a first and important step towards a global herbarium. Now, the entire collection of occurrences can be used, increasing the scale of data available for a multitude of research questions, from increased access to the entire biodiversity and abundances of the collection, to focussing on occurrences of specific species which are now reachable over many collections. Through our stages of manual review and application of taxonomic knowledge, our workflow has recorded all decisions made concerning nomenclatural corrections and the removal of names (Discussion S1), allowing for a transparent process which can be updated as new occurrences are added to the metacollection.

The process of flagging and removing names was a critical component of the curation process. There were several accepted names of flagged taxa which did not reasonably belong to the flora of Greenland, but which were only discovered and removed by checking their associated media files and related metadata after they were flagged. For example, there was one occurrence of *Arthropteris tenella* (G.Forst.) J.Sm., (GBIF catalogue number US 1371896) as a flagged name which was present in the Taxon List, which upon investigation, stemmed from an herbarium sheet with a specimen from Brisbane, Queensland, Australia. However, the word “Queensland” looks like “Greenland” due to the lettering of the handwriting, and so this specimen has been catalogued as Greenlandic.

As with any collection, there are taxa which will be rare, newly collected, or invasive, and the flagging of their names can bring their identity to the attention of users who are seeking to access this information within the metacollection. Taxonomic expertise and a knowledge of the local flora can be used to determine if such cases should be unflagged or corrected and returned to the Taxon List. For example, *Lupinus nootkatensis* Donn ex Sims (a nitrogen fixing legume) was introduced to Greenland in the 1970s (Magnusson, 2010). It is a species known to rapidly spread in other Arctic regions, such as Iceland, with widespread impacts on local diversity (Benediktsson, 2015). In our Taxon List, this species was flagged. Using our own knowledge of this species’ presence in Greenland and finding that some of the human observations were made by experts of the Greenlandic flora, we were able to confidently unflag it and keep it in the Taxon List.

The Taxon List we present is not a final nor definitive list of Greenland’s flora. Given that it is a summary of the occurrence data available, integrated with the Panarctic Flora, it is a first edition of an iterative nomenclature harmonisation and overall curation process. As existing data continues to be reviewed and cleaned, and as new data is uploaded, the inclusion of names within the collection will be further refined. While there are limitations to this workflow (Discussion S2), our current Taxon List is now the most comprehensive and accurate to date.

### Leveraging the metacollection unveils increased species diversity

The comparison of our Taxon List to the five external lists led to an increase in species. By generating the curated metacollection, we were able to create a comprehensive summary of all accessible historic collection efforts to describe the biodiversity of Greenland, stemming from many institutions and over centuries of collecting. This has allowed for a previously unprecedented scale of data to be leveraged. Greenland includes all Arctic bioclimatic zones (Raynolds et al., 2019), resulting in a wide diversity of ecological niches available to species (Bay, 2020). With this consideration, it has been acknowledged that the reported vascular plant diversity of Greenland is lower than could be expected (Bay, 2020), given the availability of such niche space and the diversity in bioclimatic zones (Bay, 2020; Normand et al., 2013). The increase in diversity revealed by the metacollection is therefore not altogether surprising, as it is an expected property not only of the Greenland environment, but also of the creation of a large and summarising metacollection.

### Nomenclature harmonisation improves taxonomic accuracy within the metacollection

A reduction in taxon richness is an expected result for any flora undergoing this workflow, along with the significant increase in the number of occurrences per accepted name. Here we demonstrate that the harmonisation process for the Greenland flora also reduces evenness within the metacollection while increasing dominance. This indicates that the harmonisation process largely reallocated GBIF occurrences to accepted names with higher numbers of occurrences within the collection, suggesting that there are certain taxa which are consistently collected (but under many names) more than others. A large portion of the occurrence data derive from vegetation surveys of Greenland in the 1980s (Bay, 2020; Bay et al., 2017; Iwanycki Ahlstrand, 2023). As such vegetation surveys will likely reveal a major proportion of the local flora, the integration of the reference flora unsurprisingly had little effect on the diversity measurements (even when looking at taxa richness).

### Unrecognised Names are of taxonomic interest

Globally, it is estimated that 70,000 flowering plant taxa are yet to be described, and that over half of them have already been collected within herbaria (Bebber et al., 2010). For many taxon names, there is a lag of several decades between discovery (and description) and the publication of that description (Bebber et al., 2010). Moreover, even when a taxonomic classification has been published, it does not mean that the publication is both digital and accessible (Pyle, 2016). This can result in taxonomic determinations, an important output from a scarce human resource, being inaccessible and unintegrated, resulting in names not being recognised (Discussion S3).

### Enhanced data availability for a region undergoing rapid change

Access to precise temporal data is critically important in understanding how climate change and other perturbations to ecosystems will impact biodiversity. A recent study on spring phenology in Greenland demonstrated that even a small increase in temporal data in regions undergoing change is critical for understanding both long-term changes in environment, and long-term responses and sensitivities of species within these environments (Schmidt et al., 2023). Given that our harmonisation process increased the average time span of occurrences in accepted names which existed in the Taxon List both before and after harmonisation by 15.7 years, the relevance in harmonising a metacollection for aiding in research that mandates strong temporal resolution is evident.

## Conclusion

Through this workflow, we have demonstrated that vast collections of occurrence data, from numerous sources and in numerous states of curation quality, can be nomenclaturally harmonised and merged by accepted name into a metacollection. The availability of this metacollection opens new research opportunities for understanding how plants are distributed through space and time, and how they respond to climate and environmental change.

## Acknowledgements

This research has been supported by the Carlsberg Foundation Semper Ardens: Accelerate grant for Natasha de Vere at The University of Copenhagen, under the title of “Greenland Plant Diversity and Pollination Networks in a Changing Arctic”. NK and GL were supported by the Swiss National Science Foundation (PZ00P3_202127) and by the Italian Ministry of University and Research (PRIN 2022 PNRR P2022N5KYJ). We would like to express our gratitude to the contributors of the Annotated Checklist of the Panarctic Flora Vascular (PAF) plants, with special appreciation to Cam Webb (University of Alaska Museum of the North) for providing an Excel version of the PAF. We are also thankful to Andrea Hahn, Head of Data Products at the Global Biodiversity Information Facility, (GBIF) for providing insights surrounding GBIF’s taxonomic backbone. We would also like to thank the many herbaria around the world which have been digitising their specimens and making them publicly available on platforms such as GBIF, allowing us to access their data and form a metacollection. This unprecedented access to digitised specimen data would not be possible without the tireless efforts of curators, botanists, volunteers, and engaged community members.

## Author Contribution

Natasha de Vere, Brandon Samuel Whitley, and Natalie Iwanycki Ahlstrand conceived and designed the research. Brandon Samuel Whitley performed the research and analysed the data with input from all authors. Brandon Samuel Whitley wrote the manuscript with input from all authors. All authors contributed to the final submitted manuscript.

## Data Availability Statement

The data that supports the findings of this study are openly available from https://www.gbif.org/ at https://doi.org/10.15468/dl.yta9wk

The script that supports the processing of this data is openly available at https://github.com/BrandonSamuelWhitley/GBIF_Occurrence_Curation

