## Supporting Information for "Harmonising digitised herbarium data to enhance biodiversity knowledge: creating an updated checklist for the flora of Greenland"

The following Supporting Information is available for this article:

**Methods S1** Detailed Workflow Methodology

**Figure S1** Partitioning of the Taxon List Accepted Names by taxon rank and data source.

**Figure S2** Specimens per Accepted Name in the pre-curation and post-curation datasets.

**Table S1** Chi-Square Test Results for Taxon Ranks

**Table S2** Adjusted pairwise comparisons from the original GBIF data.

**Discussion S1** – Removing Names

**Discussion S2** – Limitations to Workflow

**Discussion S3** – Unrecognized Names of taxonomic interest

**Dataset S1** – Workflow Outputs - Names

**Dataset S2** – Workflow Outputs - Occurrences

**Methods S1** Detailed Workflow Methodology

*Selection of Data Source*

GBIF was selected as the primary data source for accessing multiple collections of digitised herbarium specimens and other occurrence data. GBIF is a globally accessible online data infrastructure which allows for biodiversity data of numerous forms to be uploaded and linked across collections and other pathways, such as by geography. To date, GBIF contains over 468,000,000 individual data points of Tracheophytes, with over 100,000,000 coming from the digitisation of preserved specimens (GBIF.org, 2024).

*Selection of Taxonomic Backbone*

Plants of the World Online (POWO) was selected as the standard taxonomic backbone off which to base the workflow. POWO was founded in 2017, and functions as a continually updated repository of plant information, which includes for a given taxon its full name (accepted spelling with author), the status of the name (accepted, synonym, or unplaced), known synonyms of the accepted name, and other associated data such as geographic distribution (POWO, 2024). POWO is based on the World Checklist of Vascular Plants, which is built around the International Plant Names Index (IPNI), an internationally recognised authority on nomenclatural data for a given taxon, including names of infraspecific ranks (Govaerts et al., 2021).

*Mass Curation Workflow*

The *rgbif* package was used to launch a GBIF query (https://doi.org/10.15468/dl.yta9wk) (Chamberlain, 2017). GBIF query filters were set to include only specimens of the Tracheophyta taxa ID (customisable), coming from Greenland (customisable for up to three countries at once), of a "present" occurrence status, and of all bases of records excluding fossil specimens. In addition to herbarium specimens this query also included human observations (customisable). Human observations are typically photographed living specimens in the field.

Initial scientific names which GBIF had altered to a higher taxonomic rank were restored to their original inputted (verbatim) scientific name. This case often applied to hybrid names (which were altered to a family level name). Other cases when GBIF altered scientific names to a higher taxonomic rank included occurrences flagged by GBIF as having names which could not be distinguished properly (often due to a lack of author information). In this case, these names were first processed by checking for available author information with the occurrence data and appending this to the taxon name when possible. Using the `KEWR` package (Walker, 2023), remaining names of this category were checked against POWO to see if there was only one author option. If only one option existed (and the name was therefore also recognised by POWO), that author information was used to update the name. If multiple author options existed and the name was recognized by POWO, these names were set aside from analysis as they could not be confidently harmonised and were stored as a component of the removed names output. All removed names were eventually stored in a Removed Names list, where each removed name is recorded along with the reason for its removal. If the name was not recognised by POWO, they were set aside for later processing as Unrecognised Names.

For the remaining occurrences, the unique GBIF interpreted scientific names were extracted (the names which become the name of the occurrence). Names above the rank of species were removed from the initial query and stored, including cases for which the specific epithet was unknown. To effectively query against the POWO database, the format of infraspecific rank markers, such as subspecies, were changed to The International Code of Nomenclature for algae, fungi, and plants (ICNafp) standards to match the required formatting of the POWO database. Author names were then removed using the `FLORA` package in R (customisable) (Carvalho, 2024) to enable query against the POWO database. The original interpreted scientific names from GBIF were stored to later match the occurrence with its verified name.

The list of names was queried against the POWO database. Query results were binned according to if they were already considered the Accepted Name for the taxon (to have an accepted nomenclatural status), if they were a synonym for the Accepted Name of the taxon, or if the name was not recognized by POWO. Synonym names were tied to an associated Accepted Name, from which a unique Taxon ID was extracted. The Taxon IDs were also extracted directly from the Accepted Names.

The compiled list of Taxon IDs was queried against POWO to extract all pertinent metadata for each Accepted Name (with author), including all recognised synonyms. Synonyms were originally packaged as lists within the dataframe but were then unlisted and combined with their author names to avoid matching (illegitimate) homonyms belonging to different species. Illegitimate homonyms are names which are repeated for multiple taxa, but which describe different taxa, and therefore are not the same thing. Taxonomically unplaced names were removed to the Removed Names list if there was an equivalent placed name available.

The original names assigned to the occurrences of the given taxa were then appended back into the list of Accepted Names and other metadata. This allowed the source data from GBIF to be connected to the harmonised data. To append this data, the original names were compared to all Accepted Names and synonym names for each taxon, maintaining author names to account for illegitimate homonyms. The list was then filtered (against the input data) to remove homonym taxa which had been accrued due to POWO queries only working without author names. Removed names and affiliated metadata were stored for manual checks. Names above the rank of species were then queried against POWO, with query results being filtered against matches to the original inputted names.

After these steps, a preliminary Taxon List containing known Accepted Names, and a list containing Unrecognised Names (Names not recognised by POWO) was produced. These Unrecognised Names required manual checking by a taxonomic expert or an individual possessing basic knowledge of plant taxonomy, nomenclature, and the interpretation of digitised records.

*Manual Taxonomic Checking – Autonyms*

Autonym names are names which contain the epithet for the typical infraspecific taxon (such as the subspecies) and therefore the same epithet as the specific epithet (e.g. *Saxifraga paniculata* subsp. *paniculata*). Autonym names unrecognised by POWO were manually checked. These cases were removed from the Unrecognised Names using a text pattern. Once removed, the spelling of the genus and specific epithet were checked against POWO to correct cases where spelling differed. Cases where the parent species of the autonym was itself a synonym resulted in the updating of the autonym to be an autonym of that Accepted Name (e.g. *Hierochloe alpina* subsp. *alpina* being changed to *Anthoxanthum monticola* subsp. *monticola* since *Hierochloe alpina* (Sw. ex Willd.) Roem. & Schult. is a synonym of *Anthoxanthum monticola* (Bigelow) Veldkamp). In cases where that Accepted Names was itself a rank below species level, the synonymy of the name took precedence and the autonym was ascribed as a synonym to that Accepted Name, without being altered. Cases where there were multiple options of species rank to which the autonym could belong (e.g. *Calamagrostis stricta* subsp. *stricta* could be the autonym of *Calamagrostis stricta* (Timm) Koeler or *Calamagrostis stricta* Hegetschw.), were removed from the workflow and placed in the Removed Names list. Remaining autonym names were added back into the Taxa List based on Article 26 of the ICN. While these names are valid, they were likely not published and were hence not recognized in the POWO backbone, even though this is recommended by ICN recommendation 26B.1.

*Manual Taxonomic Checking – Remaining Unrecognised Names*

Remaining Unrecognised Names were checked, whereby for each name, one or more occurrences and their digitised records were examined by accessing the media file associated with the GBIF record, such as a digitised herbarium sheet. We then manually confirmed whether the taxon name and/or author provided to GBIF differed from that on the actual label (a transcription error), and whether there were any other indicators with the occurrence to justify the name not being recognised. During this manual check stage, POWO Accepted Names were linked to the Unrecognised Names whenever possible. Unrecognised Names could be described in five main categories: 1. Names with minor differences in spelling to the taxon name, which could be corrected 2. Names with minor differences in spelling to the author, which could be corrected. 3. Names where the specimen's label differed from the GBIF interpreted scientific name, and this name was recognised by POWO, which could be corrected. 4. Names which were not recognised by POWO, but which may be specific to the location of query and thus needed to be checked in another more localised reference flora. These names needed additional corroboration. 5. Names where the specimen's label differed than the GBIF interpreted scientific name, but this name was not recognized by POWO. These names needed additional corroboration.

Names which could be corrected were done so manually. After corrected names were updated, they were first searched against matches already found (Accepted Name and synonyms), in case they were corrected to an existing Accepted Name in the Taxon List. Remaining names were then queried against the existing removed illegal homonym downloads from POWO, to reduce computation time. Thereafter, any remaining names were freshly queried against POWO.

The preliminary Taxon List containing known Accepted Names was updated to account for corrected names, and the list containing Unrecognised Names (Names not recognized by POWO) was updated to remove corrected names.

*Selection and integration of an external reference flora*

An external reference flora more localised to the geographic region of focus was selected. Criteria for selection included that the flora had to be digitally available, recently updated (or able to be updated via contacting a listed editor), and that the flora is considered to be comprehensive for the scope of the workflow. For this case study of Greenland, the Annotated Checklist of the Panarctic Flora Vascular plants (PAF) was selected (Elven et al., 2017). This flora is integrated into the Taxon List, such that localised taxonomic knowledge not found on POWO could be secured, and to enable comparisons between the two data sources.

As an initial integration step, the remaining Unrecognised Names were queried against the PAF reference flora (Accepted Names and synonyms). Any matches were removed from the Unrecognised Names list. The final remaining Unrecognised Names, containing names of taxa not found in the POWO backbone nor the localised reference flora, was amalgamated into the Removed Names list.

To integrate the two data sources, the preliminary Taxon List and the reference flora were queried against one another across both Accepted Names and synonyms. To automate the workflow as much as possible, perfectly matching names were connected, ignoring differences in accents and spacings. This also enabled matching to occur to more than one preliminary Taxa List Accepted Name (indicating that the given reference flora Accepted Name matched to more than one POWO Accepted Name). Thereafter, a fuzzy join of 90% similarity was used to suggest imperfect matches between the preliminary Taxon List and the reference flora. These suggested matches were manually checked, using knowledge of the local flora to determine integration. All Accepted Names and synonyms in the reference flora were individually matched to ensure all double matches were captured.

Matches were examined per Accepted Name in the reference flora. All correct matches were linked using the POWO Accepted Name as the true Accepted Name, and appending in the reference flora names as synonyms. The cases where an Accepted Name or synonym(s) of the reference flora matched to more than one POWO Accepted Name were removed and appended to the Removed Names list.

After integration, the Taxon List contained Accepted Names derived from both the GBIF data and the POWO backbone, as well as the local reference flora. All Accepted Names in the Taxon List were then allocated a data source value to indicate if a taxon was found in the GBIF data, the reference flora, or both. If a reference flora name was removed for having multiple POWO matches, it was not used in the allocation of data source due to different taxonomic barriers existing between the POWO backbone and the reference flora.

*Data recovery – processing Unauthored Names*

The remaining Unauthored Names were queried against the Taxon List. When there was a case of an Unauthored Name matching to one Accepted Name or synonym, and there was only one possible match in the dataset, then the Unauthored Name information was appended with the appropriate author designation, given that this taxon was therefore known to occur in this dataset. Remaining Unauthored Names were appended into the Removed Names list. The same process was used for the removed unauthored names which had multiple author options, whereby if one of the multiple options was found in the Taxon List (but not more than one), then it was appended to that taxon.

*Internal validations – flagging names of concern*

Using the final Taxon List, the GBIF data was appended with the final harmonised Accepted Name for the given specimen, based on the workflow. A single taxon of a given Accepted Name could now be queried across all the GBIF data, and all occurrences matching to that Accepted Name or any of its synonyms could be gathered.

To internally validate the data, this step was performed on each Accepted Name. In rare cases where a synonym was shared across two or more Accepted Names (whereby the POWO backbone listed the name and author as a synonym to more than one Accepted Name), then the GBIF occurrence was labelled with all Accepted Names separated by "or", to indicate that a single Accepted Name cannot be allocated (e.g. *Cerastium vulgatum* L. is a synonym of both *Cerastium glomeratum* subsp. *glomeratum* and *Cerastium holosteoides* Fr.).

Validation criteria was set such that any Accepted Name containing five occurrences or less (customisable), or any Accepted Name only based on human observation (and not digitised specimens) was flagged. Additionally, an option was coded to flag names based on a minimum number of contributing data sources (customisable), to enable further customisation for other users. Autonyms were excluded from flagging, given their automated generation and validity.

*Internal validations – applying taxonomic expertise*

The resulting Flagged Names contained names which were either rare taxa in the collection, unpublished determinations, incorrect determinations, and/or containing other issues such as specimens with Accepted Names which were incorrectly geographically assigned to Greenland. We applied an automated and manual process to determine if these Flagged Names needed to remain flagged, be corrected, be unflagged, or be marked as having low data quality. To first automate as many cases as possible, names were unflagged when their data source included that of the reference flora and when the POWO backbone recognised the name as occurring in Greenland (customisable). Moreover, flagged names of the Genera rank or higher were unflagged if a taxon within their rank at a lower rank level was in the list and not flagged, giving validity to the presence of the higher taxon rank levels.

Remaining flagged names were checked, applying taxonomic expertise to determine the outcome of the check. For the Flora of Greenland, we first unflagged cases where human observations were made by known experts in the field. We then unflagged names that were found in external databases of Greenlandic plants. Thereafter, we checked occurrences for each individual flagged name. We unflagged cases when there was clear evidence that the name belonged to the flora (multiple independent data sources, collection under multiple synonyms, visual digitised specimen verification using expertise on the local flora, evidence for collection within Greenland, DNA barcode confirmation, and confirmed external verifications by other taxonomic experts). When digitised specimens were found to be incorrectly transcribed, we corrected their name. When digitised specimens were found to contain illegible labelling, or if they contained other properties whereupon we had no confidence in the name, we marked them as having low data quality. When a name did not meet criteria for these other labels, it was left as flagged.

*Internal validations – cleaning the Taxa List*

After reviewing the Flagged Names, any Accepted Names marked as having low data quality were removed from the Taxon List and appended into the Removed Names list. Remaining Flagged

Names also underwent this process. Unflagged Accepted Names were left, while Accepted Names found to be improperly transcribed were removed from the Taxon List. The updated names from these cases were then checked against the existing Accepted Names and their synonyms. If no match was found, these were freshly queried against POWO. When they were allocated to an Accepted Name, their number of occurrences was again checked against the customised criteria to determine if they were removed or not from the Taxon List. If removed, these cases were appended into the Removed Names list, along with updated names not recognised (including synonyms found in the GBIF occurrence data).

*Comparison to External Floras*

The remaining Taxon List was then compared to external floras for Greenland. This part is customised to Greenland, however for all users, the default external list is that of the KEW POWO database, which is automatically generated for the countries being queried. The user then has the option to supplement this list or use it on its own. For Greenland, the Taxon List was compared to five external floras:

(1) The Greenland Flora (1978, updated in 2020) – a checklist of native and naturalized taxa listed in the 1978 Grønlands Flora (Böcher et al., 1978), transcribed and updated in 2020 (Bay, 2020; Jacobsen et al., 2020).

(2) The 2011 Wildflowers of Greenland – a field guide for the vascular plants of Greenland (Rune, 2011).

(3) The KEW POWO flora listed for Greenland – flora listed as occurring in Greenland in the KEW POWO database (POWO, 2024)

(4) The 2018 Red List Assessment for Greenland's Plant Species (Boertmann & Bay, 2018).

(5) The 1972 Adventitious plants and cultivated plants in Greenland – a checklist of introduced plants found in Greenland between 1806 and 1972, transcribed in 2023 (Jacobsen, 2023; Pedersen, 1972).

Using these external lists, the Taxon List was compared and then species from the Taxon List not found at species level or any other infraspecific rank level in the external lists were isolated for manual review. These species unique to the Taxon List were checked and validated using the images attached to the occurrences, knowledge of the flora, and individual research on each species. Species were removed from the Taxon List following the same removal protocols as the flagged names, and if removed, were added to the Removed Names list.

*A Final Taxa List*

After processing all Flagged Names, the final Taxa List was produced, where each remaining Accepted Name could be confidently included in our flora of Greenland checklist. As an additional output option depending on user needs, names above the rank of species could be kept only when there were no cases of a species within that higher rank being present in the dataset. Additional outputs of the process included the Removed Names list, containing all names that had been removed during the workflow (along with their taxon ranks, reasoning about why they were removed), and Specimen Counts data, where for both the Taxon List and the Removed Names list, the number of occurrences per Accepted Name were tallied, as well as their age-range. Finally, two GBIF occurrence files were produced, one for the Taxon List and one for the Removed Names list, where every occurrence was assigned the Accepted Name or the equivalent, allowing for the occurrence data to be treated as one metacollection.

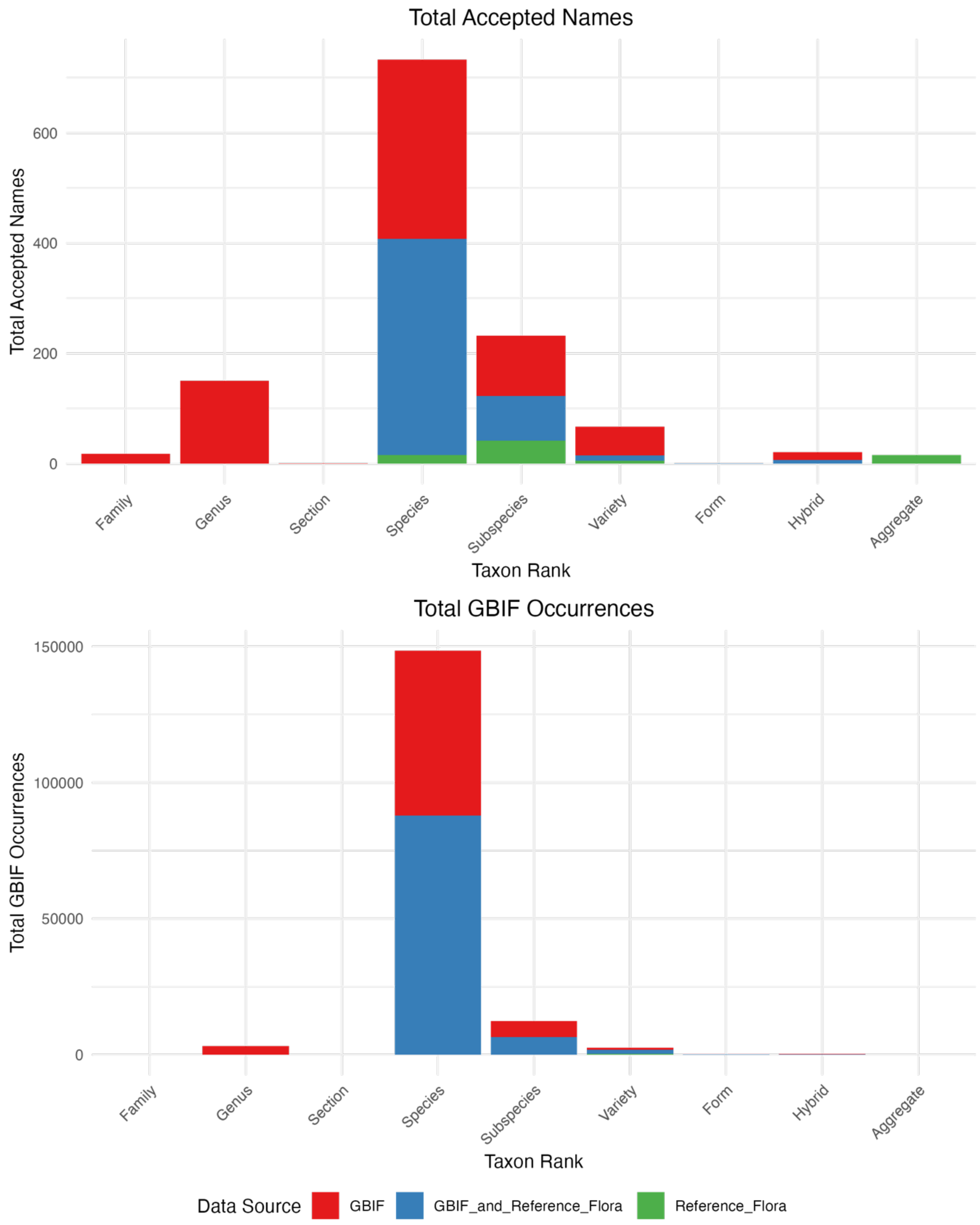


**Fig. S1** Partitioning of the Taxon List Accepted Names by taxon rank and data source, with matching distributions of the affiliated GBIF occurrence data. Species-level taxa dominate both the Taxon List and the affiliated occurrences.

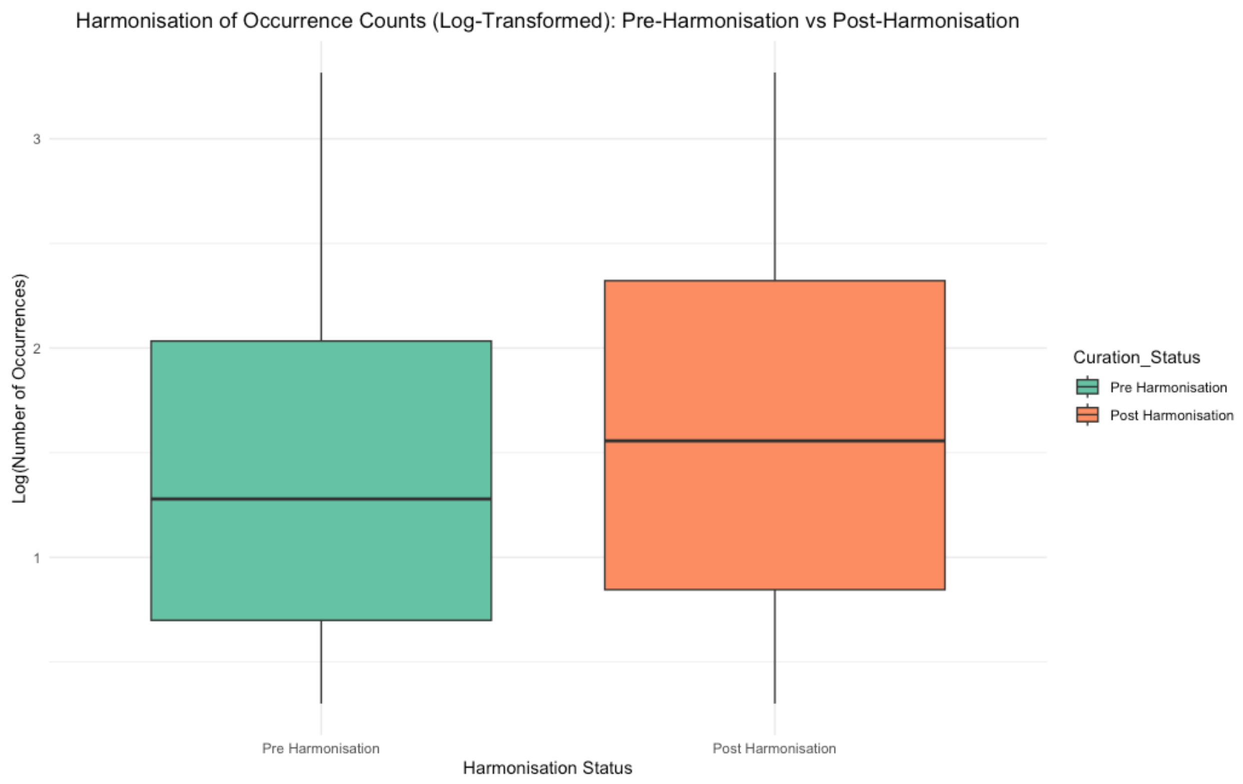


**Fig. S2** There is a significant difference between the number of specimens per Accepted Name in the pre-curation and post-curation datasets (Wilcoxon signed rank test with continuity correction, $V = 160472$, p-value = 6.558e-07). Data was not normally distributed (Shapiro-Wilk normality test, $W = 0.85643$, p-value < 2.2e-16). Note that the comparison was only made between Accepted Names shared by both datasets.

**Table S1** Chi-Square Test Results for Taxon Ranks, indicating that in almost all cases there was a significant decrease in alpha diversity of the GBIF occurrence data during harmonisation.

| Taxon Rank | X-squared | Degrees of Freedom | p-value | Significance |
|---|---|---|---|---|
| Form | 214.16 | 2 | < 2.2e-16 | Significant |
| Hybrid | 3.2703 | 2 | 0.1949 | Not Significant |
| Species | 396.43 | 2 | < 2.2e-16 | Significant |
| Subspecies | 70.813 | 2 | 4.199e-16 | Significant |
| Variety | 583.23 | 2 | < 2.2e-16 | Significant |

**Table S2** Adjusted pairwise comparisons using Fisher's Exact Test with Bonferroni correction from the original GBIF data (Pre-Harmonisation) to the curated GBIF Data (Post-Harmonisation) to the final harmonised data integrated with the Panarctic Flora (Integration of Reference Flora).

| Pairwise Comparison | p_value | adjusted_p_value | Significance |
|---|---|---|---|
| Species: Pre-Harmonisation vs. Post-Harmonisation | 4.1e-97 | 6.2e-96 | Significant |
| Species: Pre-Harmonisation vs. Integration of Reference Flora | 7.4e-93 | 1.1e-91 | Significant |
| Species: Post-Harmonisation vs. Integration of Reference Flora | 6.5e-01 | 1.0e+00 | Not Significant |
| Subspecies: Pre-Harmonisation vs. Post-Harmonisation | 3.0e-22 | 4.5e-21 | Significant |
| Subspecies: Pre-Harmonisation vs. Integration of Reference Flora | 2.0e-13 | 3.0e-12 | Significant |
| Subspecies: Post-Harmonisation vs. Integration of Reference Flora | 2.0e-02 | 3.0e-01 | Not Significant |
| Form: Pre-Harmonisation vs. Post-Harmonisation | 2.5e-61 | 3.8e-60 | Significant |
| Form: Pre-Harmonisation vs. Integration of Reference Flora | 2.5e-61 | 3.8e-60 | Significant |

| | | | |
|---|---|---|---|
| Form: Post-Harmonisation vs. Integration of Reference Flora | 1.0e+00 | 1.0e+00 | Not Significant |
| Variety: Pre-Harmonisation vs. Post-Harmonisation | 2.0e-145 | 3.0e-144 | Significant |
| Variety: Pre-Harmonisation vs. Integration of Reference Flora | 3.2e-141 | 4.7e-140 | Significant |
| Variety: Post-Harmonisation vs. Integration of Reference Flora | 7.1e-01 | 1.0e+00 | Not Significant |
| Hybrid: Pre-Harmonisation vs. Post-Harmonisation | 8.6e-02 1.0e+00 | 1.0e+00 | Not Significant |
| Hybrid: Pre-Harmonisation vs. Integration of Reference Flora | 8.6e-02 | 1.0e+00 | Not Significant |
| Hybrid: Post-Harmonisation vs. Integration of Reference Flora | 1.0e+00 | 1.0e+00 | Not Significant |

**Discussion S1** – Removing Names

The removal of names from the taxon list acts as an important marker in the iterative curation process of which this workflow is a part. By linking the removed names back to the occurrence data, the institutions which have contributed such occurrences can be contacted regarding their data and can be notified if their occurrence data needs further validation. Once validated, the data may be corrected, which will in turn alter the taxon list in a later iteration of this process. The process of flagging Accepted Names was critical to ensuring a curated Taxon list, for this enabled cases to be checked when the name of a taxon was nomenclaturally valid but its inclusion in the Taxon List was suspect. By setting our own criteria to flag a name (customisable in our workflow), we were able to control our level of confidence in the occurrence data based on our knowledge of the data sources and quality.

**Discussion S2** – Limitations to Workflow

It is important to note that since this workflow produces a summary of occurrence data contained in the historical collections for a given region, the accuracy of the Taxon List it produces is only as strong as the representativeness of the available occurrences. With this in mind, we strongly support the herbarium specimen digitisation revolution currently underway. With each digitised specimen, more data become available for harmonisation into an increasingly powerful and representative metacollection. This process is necessary to unlock the vast research and conservation potential for the natural and social sciences alike. It should also be considered that the accuracy of the Taxon List is subject to the quality of the inputted GBIF data (correct taxa assignments), in conjunction with the name flagging criteria determined by the user. Therefore, we also advise refining the flagging criteria over several iterations to determine an appropriate threshold value given the data at hand. Finally, we note that genuine hybrids are often found to be unrecognised in the POWO taxonomic backbone, either due to their spelling, the order of the hybridised epithet names, or simply because that they are only known locally to the region. In regions which are known to have high rates of hybridisation, such as the Arctic (Abbott & Brochmann, 2003; Brochmann & Brysting, 2008), we advise checking the hybrids which were removed for being unrecognised, and to apply local knowledge of the flora to determine if they are in fact legitimate and should be re-added to the Taxon List.

**Discussion S3** – Unrecognised Names of Taxonomic Interest

During manual review stages, evaluating the Unrecognised Names and their digitised occurrence data allows areas of research interest to be noted. As an example, the genus *Ranunculus* contributed the most Unrecognised Names to the output (59 names). Upon checking the corresponding digitised herbarium specimens, it became clear that these names underwent a taxonomic review in 1982 by two taxonomists, likely experts on this genus. However, these determinations are not available digitally, so the names did not become part of the taxonomic backbone used for our harmonisation process. With the partitioning of the GBIF data into Unrecognised Names, cases like these can be noted and potentially resolved, which in turn would feed back into the iterative curation loop of this workflow.

**Dataset S1 – Workflow Outputs - Names**

A separate Excel worksheet titled "All_Taxa_List" contains the following sheets.

**Taxon_List**: Taxa found in the final curated Taxon List presented in the paper along with associated metadata, including synonyms for the Accepted Name.

**Taxon_List_Occurrence_Counts**: the number of occurrences per each Accepted Name found in the GBIF data set, and the spread of those occurrences over time.

**Final_Removed_Names_List**: the names of occurrences which were removed from the Final Taxon List, and the reasoning why.

**Final_Removed_Names_Occ_Counts**: the number of occurrences per each Verified unrecognised name found in the GBIF data set, and the spread of these occurrences over time

**New_Species_Taxon_List**: the list of species in the Taxon_List which were unique to the Taxon_List when comparing to external sources.

**Taxa only from external lists**: the 73 taxa from the 5 external lists not found in our Taxon List, along with their taxon rank and their data source.

**Dataset S2 – Workflow Outputs - Occurrences**

A separate Excel worksheet titled "All_GBIF_Data" contains two sheets.

**GBIF_Occurrences_Taxa_List**: all the GBIF occurrence data which was matched to an Accepted Name found in the final Taxon List for this Greenland case study.

**GBIF_Occurrences_Removed_Names**: all the GBIF occurrence data which was matched to a Removed Name found in the Removed Names list for this Greenland case study.
